# Mechanism of circZNF827-mediated transcriptional repression during neuronal differentiation

**DOI:** 10.64898/2026.02.17.706266

**Authors:** Ivan Zaporozhchenko, Anne Kruse Hollensen, Christian Kroun Damgaard

## Abstract

Circular RNAs (circRNAs) originate from backsplicing of numerous genes in animals, but the functions of most circRNAs remain elusive. We previously demonstrated that circZNF827 forms a complex with hnRNPL/K and its host gene-encoded protein ZNF827 that acts in the nucleus to transcriptionally repress the nerve growth factor receptor (NGFR/p75NTR) gene during neuronal differentiation (Hollensen, 2020) [1]. To explore the mechanism of action, and to assess a potential role of the circZNF827-hnRNP complex on additional loci, we scrutinized the genome-wide consequences of circZNF827 and/or hnRNPL knockdown at the transcriptomic and epigenetic level. RNA-sequencing and CUT&RUN confirmed that NGFR and additional loci are transcriptionally repressed by the circZNF827-protein complex, and that these are primarily enriched for H3K27me3 signatures. Only a fraction of the massive transcriptomic changes could be ascribed to a direct circZNF827 transcription-regulated phenotype, suggesting that initial key regulatory events elicited by the circZNF827-hnRNP complex likely lead to a secondary response, which further augments neuronal differentiation.

## INTRODUCTION

Circular RNAs (circRNA) arise from head-to-tail splicing events on predominantly protein coding transcripts through a process known as backsplicing (reviewed in [2]). CircRNA expression displays cell- and tissue-specific patterns, yet little is known about their biological function or significance. The brain is especially enriched in circular transcripts, raising the possibility of a role in neuronal development and neural function [3].

Previously we have demonstrated that circZNF827, highly expressed in differentiating neuronal cells, nucleates a complex containing hnRNPL/K and circRNA-encoded host protein ZNF827, that acts in the nucleus to transcriptionally repress the nerve growth factor receptor (NGFR/p75^NTR^) gene, linking it to neuronal differentiation and neuronal development [1].

In this study, we have investigated the epigenetic and transcriptomic effects of the circZNF827-hnRNP complex. Using Cleavage Under Targets and Release Using Nuclease (CUT&RUN) [4] we have profiled genome-wide levels of activating (H3K4me3, H3K27ac) and repressive (H3K27me3) histone modifications paired with transcriptome sequencing upon depletion of the circZNF827-hnRNP complex (Figure 1). Our results reveal that the transcriptional output from the NGFR gene is repressed by the circZNF827-hnRNP complex mainly through the addition of H3K27me3 signatures. Interestingly, these marks are often found as bivalent conformations containing also activity mark H3K4me3, which suggests a repressed state that is primed for immediate transcriptional activation. Finally, our results highlight numerous additional circZNF827-mediated effects on transcription in the context of neuronal development.

**Figure 1.**
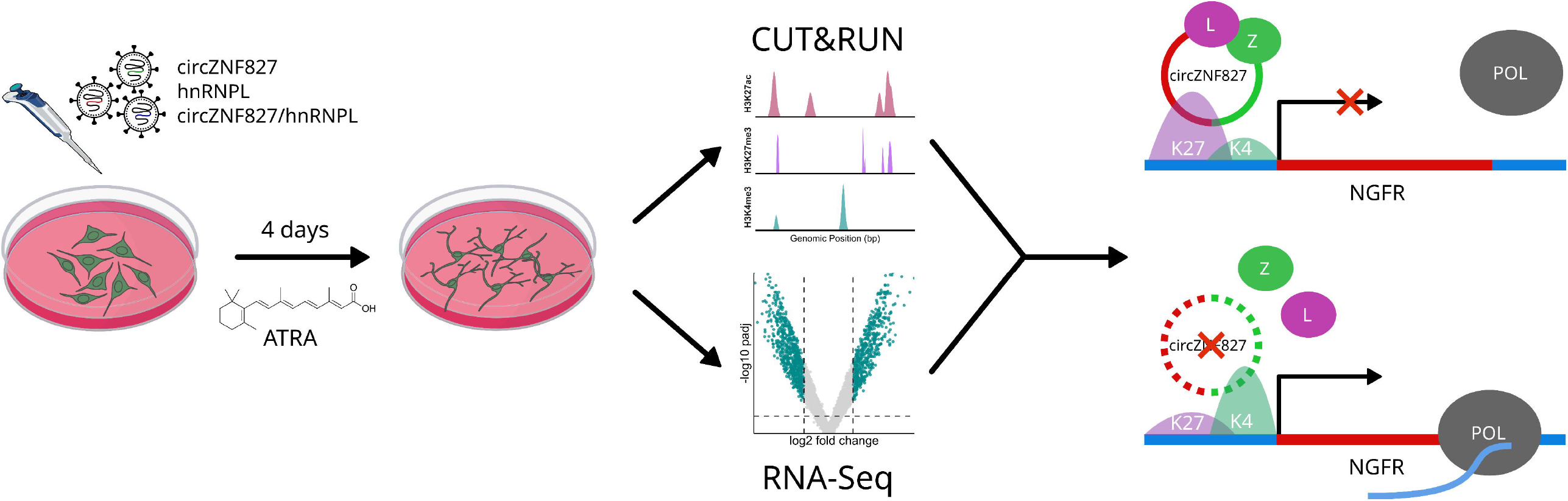
Overview of experimental strategy. Schematic illustration of the experimental workflow for lentiviral transduction, L-AN-5 differentiation, CUT&RUN and RNA-sequencing, and analysis outcomes with NGFR locus as an example. H3K4me3 - H3 lysine 4 trimethylation (K4), H3K27me3 - H3 lysine 27 trimethylation (K27), H3K27ac - lysine 27 acetylation. hnRNPL (L) and ZNF827 (Z) proteins are indicated in green and purple.

## RESULTS AND DISCUSSION

### Transcriptomic changes upon circZNF827 knockdown

Using NanoString analyses during neuronal differentiation, we have previously identified several differentially regulated transcripts upon circZNF827 depletion among ∼800 neuropathology-related genes [1]. To provide a genome-wide resolution of circZNF827-regulated genes, we subjected differentiated (retinoic acid-treated) L-AN-5 cells to knockdown of circZNF827, hnRNPL or a combined circZNF827/hnRNPL knockdown. Knockdown of circZNF827 triggered widespread transcriptomic changes (Figure 2a,b) (5363 genes, pAdj<0.05, 366 genes with abs(log2FC) > 1). Changes in transcript abundances were also detected upon hnRNPL knockdown, albeit to a lesser extent (2299 genes, pAdj<0.05, 145 genes with abs(log2FC) > 1). Approximately 27% of differentially expressed genes (DEG) after circZNF827 knockdown was also affected by hnRNPL knockdown (Figure 2c). Double knockdown of circZNF827 and hnRNPL resulted in deregulation of a unique set of differentially expressed genes (564 genes, pAdj<0.05, 62 genes with abs(log2FC) > 1). Principal component analysis identified all three knockdown conditions as well as control cells as four distinct groups (Figure 2d).

**Figure 2.**
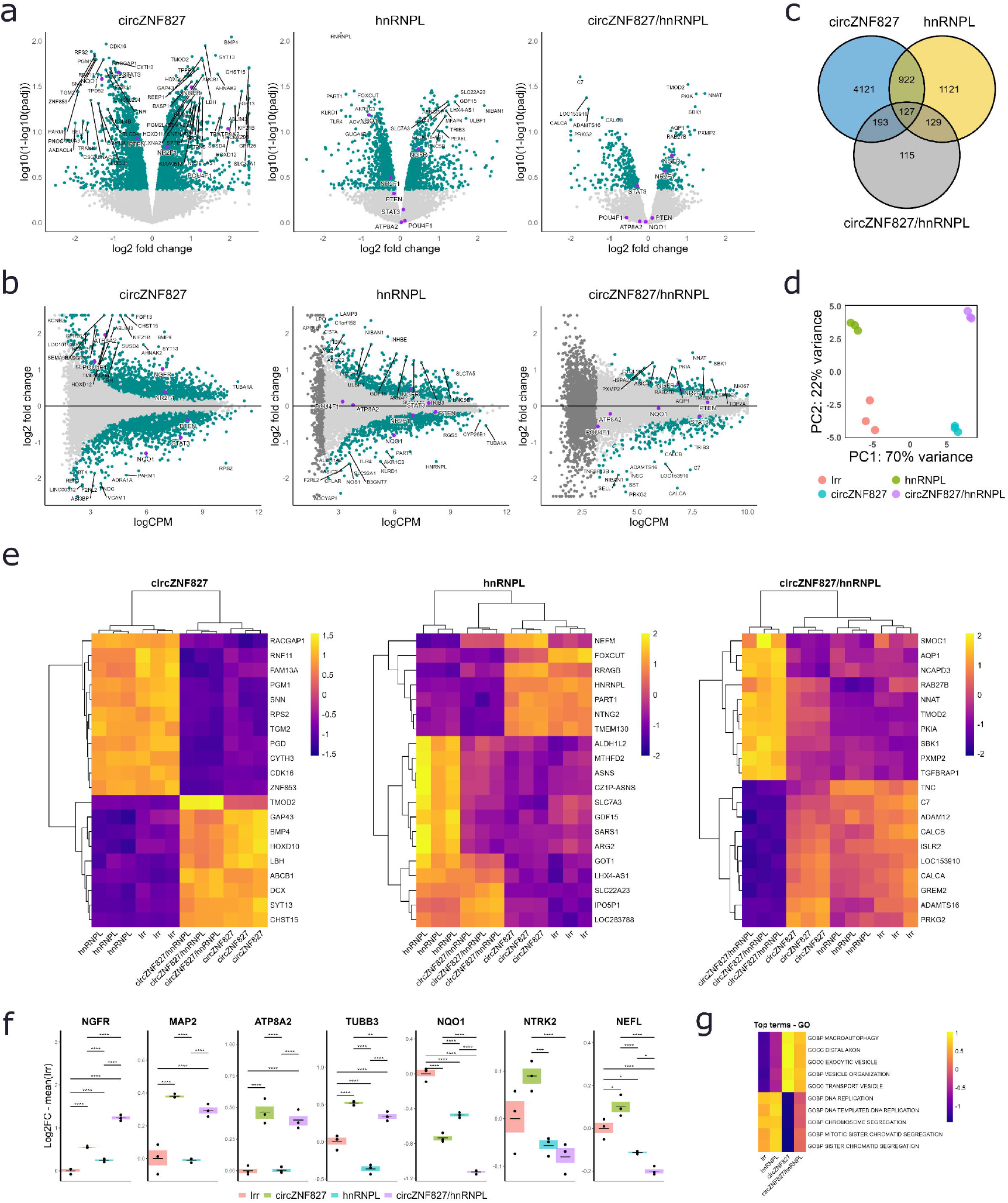
Widespread transcriptome changes identified by RNA-Seq after circZNF827 and/or hnRNPL knockdown. **(a)** Volcano plots of differentially expressed genes for three knockdown conditions. The vertical (significance) axis is transformed to log10(1-log10(pAdj)) for better clarity. (**b**) MA plots of differentially expressed genes for the three knockdown conditions. (**c**) Venn diagram of differentially expressed genes between the experimental conditions. (**d**) PCA plot of RNA-Seq samples. (**e**) Heatmaps showing top 20 differentially expressed genes across all conditions. (**f**) Boxplots showing expression of select genes. Gene expression is normalized to mean expression in control knockdown (Irr). (**g**) Heatmap of gene expression across the Gene Ontology terms enriched in up- and downregulated differentially expressed genes.

Notably, the expression of key genes and neuronal markers, including NGFR, NQO1, MAP2, ATP8A2, TUBB3 among others, known to be affected by circZNF827 knockdown, was consistent with our previously reported data (Figure 2f). In some cases, the effect of double knockdown was additive (e.g. NGFR and NQO1), while for some others (e.g. NTRK2 and NEFL) knocking down circZNF827 and hnRNPL had opposite effects on transcript abundance that were attenuated in the double knockdown. The finding that hnRNPL and circZNF827 knockdown affect primarily different subsets of genes, suggests that the circZNF827-hnRNP-ZNF827 complex could act independently of known hnRNPL involvement in regulation of multiple nuclear events, including pre-mRNA splicing/polyadenylation and general RNP maturation/export [5].

Enrichment analysis showed that the changes span beyond just neuropathology-related genes - upregulated transcripts are enriched in nervous system and neuronal component Gene Ontology (GO) terms, while downregulated were rich in cell cycle, DNA repair and replication (Figure 2g). In terms of cellular identity, transcripts common for several neuronal/brain-derived cell types are enriched among upregulated transcripts (Supplementary figure 1). Taken together, these findings are consistent with an enhanced neuronal differentiation phenotype, pushing L-AN-5 cells into a terminally differentiated G0 state upon circZNF827 knockdown.

### Epigenetic state of NGFR locus

Previously we have shown that in case of a subset of genes, including NGFR, gene expression changes caused by circZNF827 knockdown are enacted at the level of transcription. To understand the epigenetic cause of these changes, we performed CUT&RUN assays to scrutinize three common histone modifications - H3 trimethylation at lysine 4 and lysine 27 (H3K4me3 and H3K27me3, activation- and repression mark, respectively) and acetylation at lysine 27 (H3K27ac, activation mark). Consistent with transcriptional repression of the NGFR locus we observed a significant H3K27me3 decrease upon circZNF827 knockdown (Figure 3a,b). In addition, there was also a clear tendency for an increase of locus activation marks (increase in H3K4me3 and H3K27ac deposition) although these did not reach significance upon single circZNF827 knockdown (Figure 3a,b). However, this was further augmented in the double circZNF827/hnRNPL knockdown, reaching high significance also for H3K4me3 and H3K27ac. Interestingly, NGFR was also one of the top genes with correlation between H3K4me3 signatures and transcript levels across all samples (Figure 3c,d). Overall, our results show that the change in chromatin modification at the NGFR locus upon circZNF827 knockdown, is likely governed by either reduced deposition, or increased removal of H3K27me3, while H3K4me3 and H3K27ac remain less affected. Normally, the NGFR locus is likely modified with dual H3K4me3 and H3K27me3, which still confers transcriptional silencing, as H3K27me3 has been reported to “dominate” H3K4me3 in stem cells [6,7]. Hence, the NGFR gene could likely be poised for rapid activation, needing only removal of H3K27me3. Such co-existence of activating and repressive marks, termed a bivalent domain, is often seen in stem cells subjected to differentiation [8].

**Figure 3.**
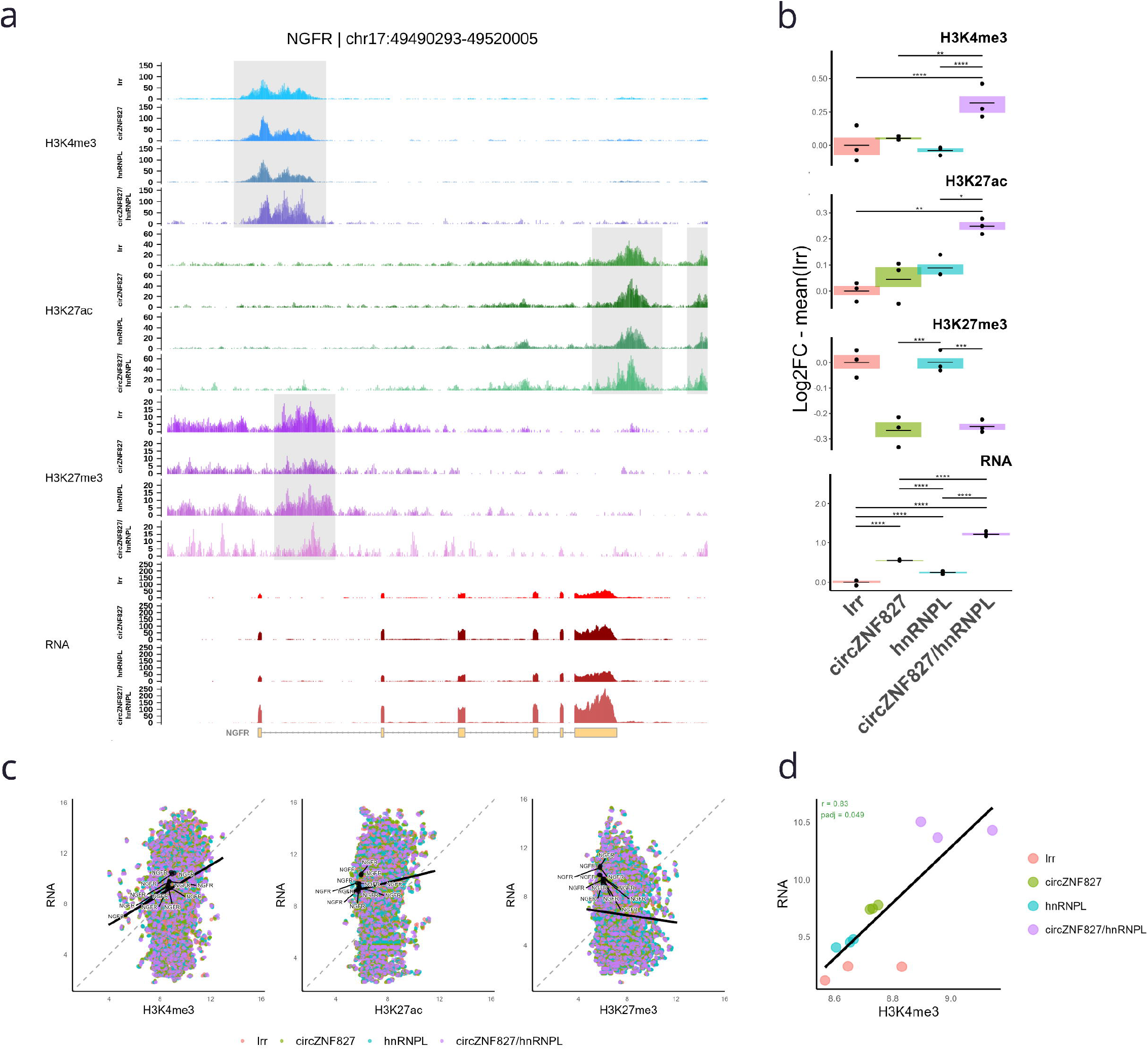
H3K4me3 and H3K27me3 signature changes at the NGFR locus after circZNF827 and/or hnRNPL knockdown. (**a**) CUT&RUN and RNA-Seq profiles. Raw data with greenlist normalization applied. (**b**) Boxplots of histone modification peaks and NGFR gene expression across conditions. Significant pairwise differences between conditions are indicated with bars. P values are reported as p<0.05 (*), p<0.01 (**), p<0.001 (***) or p<0.0001 (****). (**c**) Correlations between each histone modification and RNA abundance. Black solid line indicates the general trend across all conditions. (**d**) Correlation between H3K4me3 at the NGFR promoter and its RNA abundance across all samples. Black solid line indicates the general trend across all samples.

### Global epigenetic changes upon circZNF827 knockdown

Overall, the distribution of histone marks across the genome followed expected patterns, with major H3K4me3 located around gene promoters, H3K27me3 peaks found around and upstream of transcription start site (TSS), and H3K27ac peaks present mostly outside of gene bodies, upstream or downstream (Figure 4a,b). Knockdown of circZNF827 resulted in genome-wide epigenetic changes affecting all three histone modifications (pAdj<0.05, 1888 peaks or 1425 genes total - Table 1), while hnRNPL knockdown was responsible for fewer instances of significant shifts in histone modifications (pAdj<0.05, 631 peaks or 458 genes total, Table 1). Interestingly, most of the detected changes to H3K4me3 and H3K27ac were attributed to circZNF827 knockdown (1685 vs. 330 peaks), while H3K27me3 was found mainly upon hnRNPL knockdown (311 vs. 78 peaks) (Figure 4d,e). Overall, most significant and substantial changes were detected for H3K4me3, matching the shift in transcriptional activity (Figure 4d,e). In contrast to RNA-seq results, no significant effect of the interaction between the circRNA and hnRNPL knockdowns was detectable for H3K4me3 or H3K27ac, while a mild interaction effect was present (Figure 4e). Together, these observations suggest that the observed hnRNPL effects on bulk gene expression are mostly elicited at the post-transcriptional level (e.g. splicing, polyadenylation or RNP assembly/export) or as downstream secondary effects of these events.

**Table 1.**
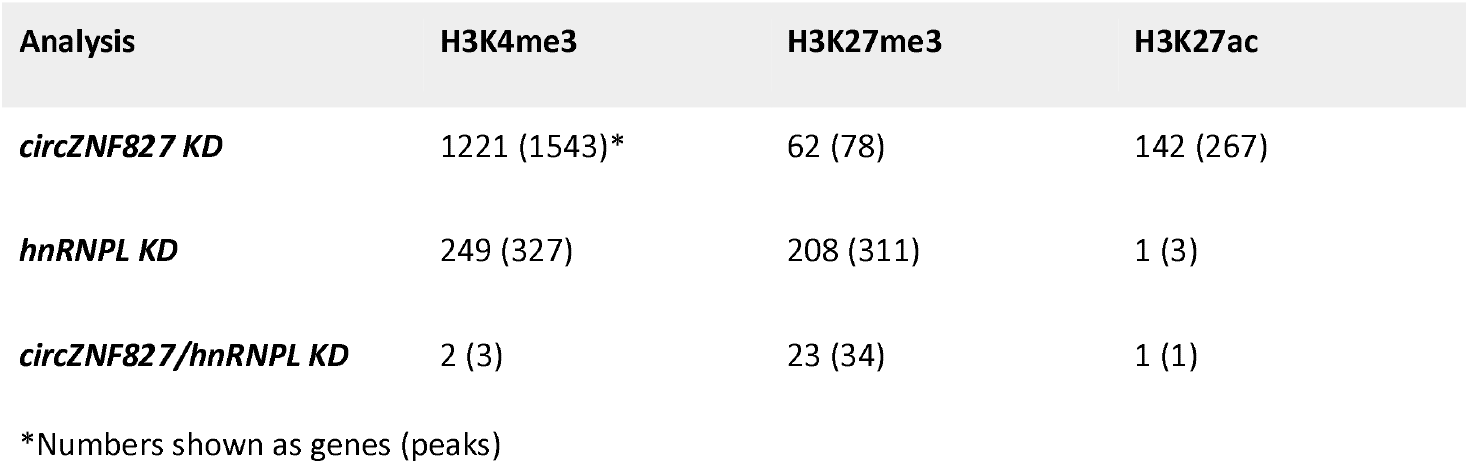
Gene and peak counts with differential intensity below significance threshold (pAdj < 0.05).

**Figure 4.**
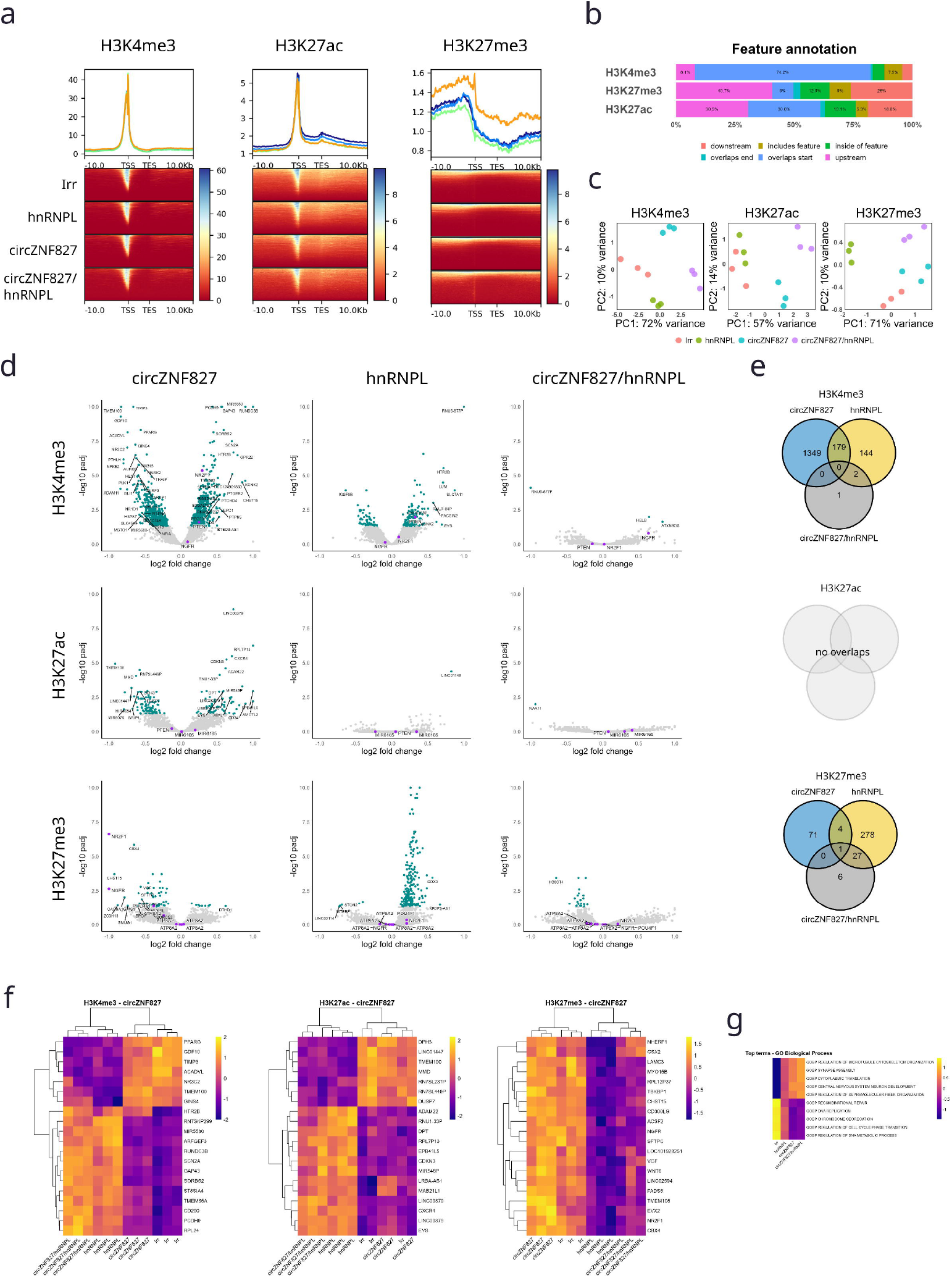
Epigenetic changes triggered by circZNF827 and/or hnRNPL knockdown are genome-wide. (**a**) Aggregated CUT&RUN read profiles within +-10 kb of known genes. (**b**) Annotation of consensus peaks to genomic features. (**c**) PCA plots of CUT&RUN samples. (**d**) Volcano plots of differentially enriched peaks for all histone modifications across the three knockdown conditions. (**e**) Venn diagrams of differentially enriched peaks between the experimental conditions. (**f**) Heatmaps showing top 20 genes with differentially enriched peaks for the three histone modifications after circZNF827 knockdown. Only peaks in proximity to known genes are shown. (**g**) Heatmap of histone modification signatures across the Gene Ontology Biological Process terms enriched in up- and downregulated differentially enriched H3K4me3 peaks.

Corroborating our transcript-level analyses, genes with increased signatures of the active chromatin mark H3K4me3, were enriched in nervous system and neuronal component GO terms (Figure 4g). In contrast, for H3K27ac, neuronal GO categories were enriched in down-regulated genes. H3K27me3 increase was primarily linked to the regulation of sodium transport. Strong cell type similarity analysis also confirmed that activated genes with increase in H3K4me3 are linked to early drivers of the neuronal phenotype (Supplementary figure 3b). No cell types were significantly enriched for H3K27me3 or H3K27ac, likely due to the lower number of DEGs/peaks below the significance threshold.

### Integrated analysis

To identify changes in gene expression directly caused by epigenetic events we compared CUT&RUN peaks and RNA-seq data. When binning H3K4me3 and H3K27ac as “active” marks and H3K27me3 as a “repressive” mark, only a fraction of genes had both significantly altered transcript abundance and a detectable change in either of the histone modifications (Figure 5a,b). Specifically, only 9% of differentially expressed genes had at least a single corresponding differentially enriched H3K4me3 peak, while for both H3K27ac and H3K27me3 modifications this number was at or below 1% (Figure 5b). A contributing factor to this may be the different sensitivity between significant calls within RNA-seq and CUT&RUN assays. However, most of the differential histone signatures did not result in significant differences in transcript production. This is supported by correlation analysis showing that only a small proportion of genes have significant correlation between epigenetic profiles and transcript abundance, most prominently for the H3K4me3 modification (Figure 3c). We conclude that most of the transcriptomic changes are likely not a direct consequence of epigenetic events related to any of the three modifications we profiled.

**Figure 5.**
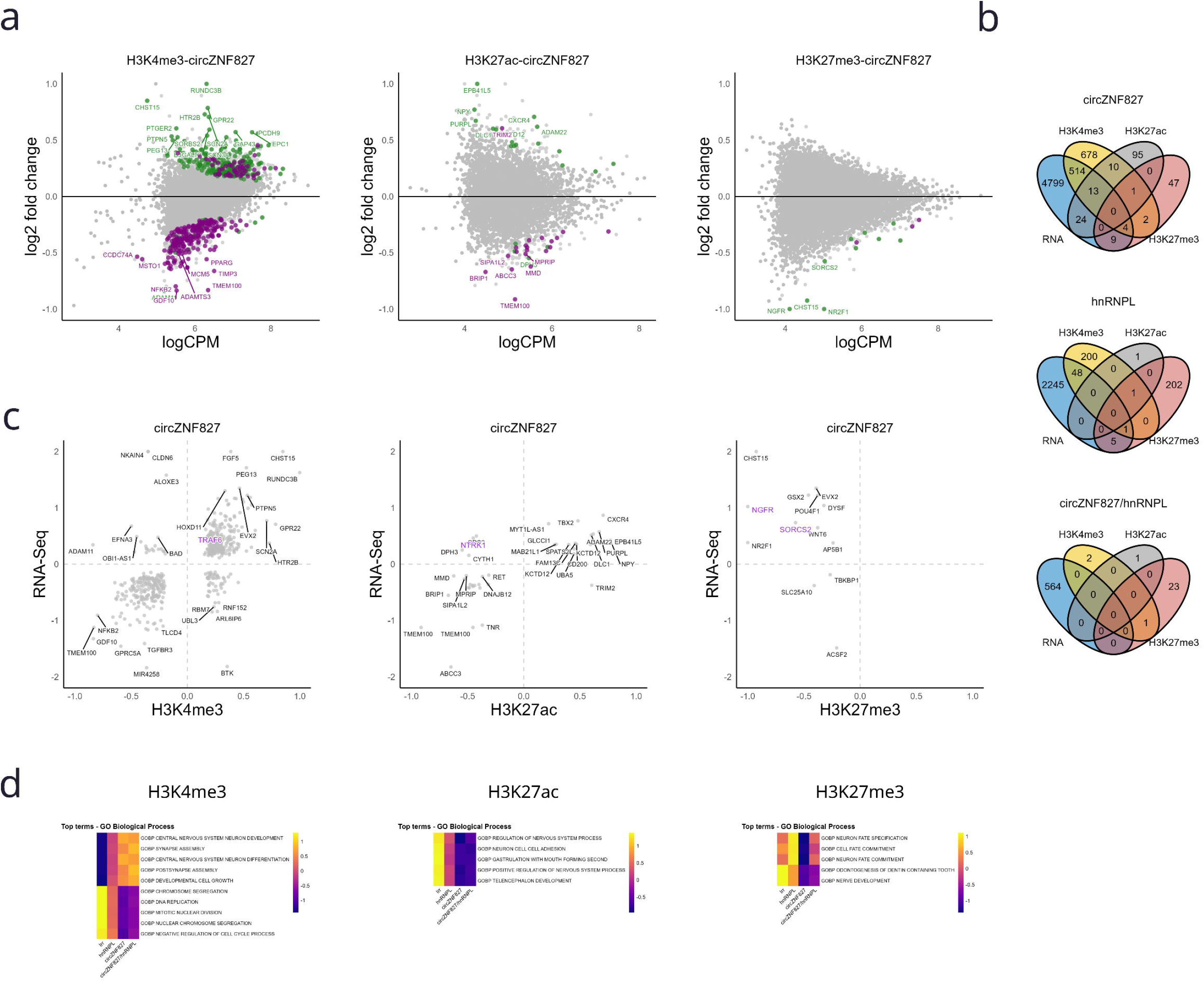
Integrated analysis highlights that only a fraction of transcriptomic change is directly caused by alteration of epigenetic profiles. (**a**) MA plots of differentially enriched peaks after circZNF827 knockdown. Genes differentially expressed after circZNF827 knockdown are labelled in green (upregulated) or purple (downregulated). (**b**) Venn diagrams showing overlaps between differentially enriched peaks of histone modifications and differential RNA abundance across the experimental conditions. (**c**) Plots showing the correspondence between changes in transcript abundance and histone modification signatures after circZNF827 knockdown. Both axes show log_2_ transformed fold change values. (**d**) Heatmaps of histone modification signatures across the Gene Ontology Biological Process terms enriched in gene sets with concordant changes in histone modification and transcript abundances.

To investigate more direct effects of circZNF827 knockdown on the epigenetic landscape, we explored the subset of overlapping DEGs and differential histone peaks that exhibited concordant direction of epigenetic and transcriptomic changes when increase in RNA output was coupled with an increase in H3K4me3 and H3K27ac peak intensity or H3K27me3 signature decrease. When focusing on this subset of genes, enrichment analysis of genes with an increase in H3K4me3 signatures showed a clear picture of neuronal differentiation in both GO terms and cell type similarity (Figure 5d). Significant enrichment of the *GO Biological Process* term *Neuronal Fate Specification* was seen for H3K27me3 decrease. For acetylation, enrichment analysis yielded neuronal differentiation terms only for a subset of genes with lower expression and decreased H3K27ac signature. Again, this supports the previously reported role of circZNF827 in “slowing down” neuronal differentiation by facilitating deposition of H3K27me3 modifications, even though we have not detected any typical H3K27me3 writers (e.g. PRC2/PRC1 complex proteins) in our previously reported circZNF827 interactome [1]. This could likely be attributed to a transient nature of a potential circRNA-protein interaction.

### Transcription factor enrichment

In order to understand the secondary effects of circZNF827 knockdown on cell phenotype, we strived to identify other potential driver genes poised for up-regulation by the decrease of H3K27me3 signatures. Among those, two transcription factors (TFs) - NR2F1 and POU4F1, were also significantly upregulated at the transcript level, while NR2F1 additionally had significantly increased H3K4me3 signature (Figure 6a). Using known binding motif sequences for both of TFs, we have ranked all possible targets and tested if they could target any genes with changes in differential expression or histone modification peak intensity. Gene Set Enrichment Analysis (GSEA) demonstrated that both up- and downregulated DEGs were enriched for NR2F1 binding sites (pAdj<0.05, Normalized enrichment scores (NES) of 1.72 and 2.07, respectively) (Figure 6c,d). This is consistent with NR2F1 being able to repress or stimulate transcription depending on promoter sequence contexts and other DNA-bound interaction partners [9]. Leading Edge gene components were subjected to GO pathway analysis showing enrichment for a variety of neuronal projection and neuronal homeostasis GO Biological Process terms for up-regulated genes and a number of *cell cycle, chromosome segregation* and *DNA replication* terms for down-regulated genes, similar to results yielded by previous enrichment analyses of these gene sets (Figure 6e). Although failing significance tests after False discovery rate (FDR) adjustment, target gene sets with decreased H3K4me3 peak intensity and lowered transcript abundance also showed similar behavior (NES = 1.68). GO enrichment analysis for Leading Edge genes of this set also showed preference for *cell cycle and chromosome segregation* terms. For POU4F1 only one gene set containing genes with decrease in H3K27ac peaks and transcript abundance levels was significantly enriched (pAdj<0.05, NES = 1.93), with GO term enrichment of Leading Edge genes showing only one statistically significant terms - *protein ufmylation*, and a number of brain development and neuronal differentiation terms just over the significance threshold (Supplementary figure 6).

**Figure 6.**
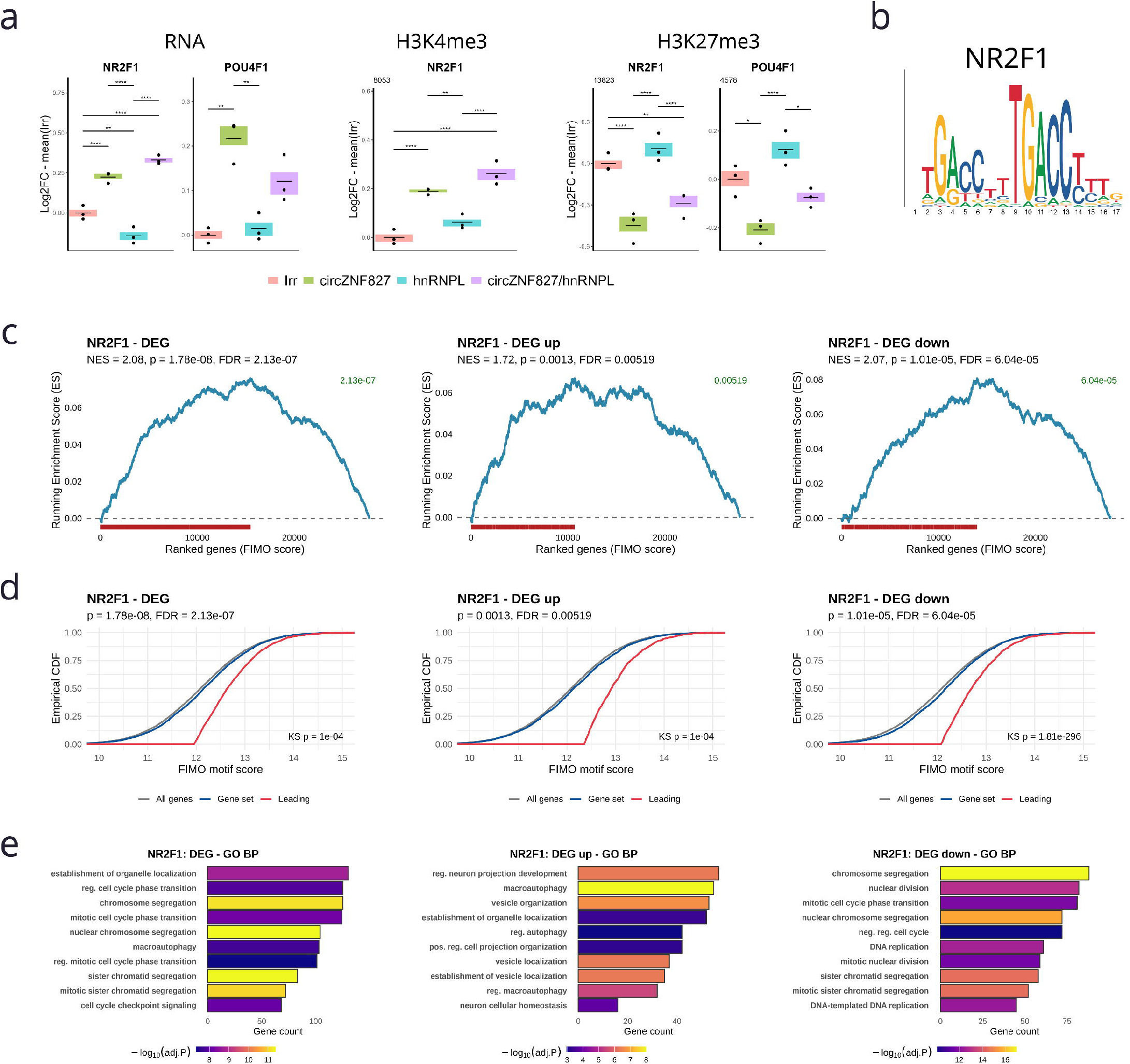
Transcription factor binding enrichment analysis identifies NR2F1 as another potential driver of neuronal differentiation. (**a**) Boxplots of histone modification abundance and gene expression of POU4F1 and NR2F1 across the experimental conditions. P values are reported as p<0.05 (*), p<0.01 (**), p<0.001 (***) or p<0.0001 (****). (**b**) Visualization of NR2F1 binding motif. (**c**) Gene Set Enrichment Analysis (GSEA) of gene sets based on CUT&RUN and/or RNA-Seq data against all predicted NR2F1 targets. (**d**) Empirical cumulative density function (CDF) plots of gene sets based on CUT&RUN and/or RNA-Seq data against all predicted NR2F1 targets. Kolmogorov-Smirnov test p-value (KS p) is given for comparison of leading edge set versus all ranked targets. (**e**) Gene Ontology Biological Process enrichments of Leading edge gene sets from the respective GSEA.

## CONCLUSION

Here we have confirmed and expanded on our previous findings that circZNF827 represses neuronal differentiation and its depletion pushes the transcriptome towards an increased output from neuron-specific genes. NGFR, known to promote either neurogenesis or cell death depending on its receptor interaction partners and availability of neurotrophins [10], is mainly affected by a decrease in the transcription repressive mark H3K27me3, potentially elicited by circZNF827-hnRNP complex-mediated recruitment of H3K27me3 writers. Strikingly, we only observe a small population of upregulated genes (RNA-seq) (266 genes, ∼1.5% of expressed genes) that displayed a similar histone methylation/acetylation profile upon knockdown of circZNF827 or in combination with hnRNPL knockdown. This finding suggests that the circZNF827-hnRNP complex only affects relatively few genes directly or that it may alter other histone signatures not tested in the present study. It remains to be investigated how the circRNA-hnRNP complex is specifically recruited to act on their repressed loci, and whether any H3K27me3 writers (PRC2-EZH1/2) become transient partners in a larger circRNA-nucleated histone-regulatory complex. The secondary output of neuronal-specific gene expression is likely facilitated by upregulation of NR2F1 or other affected transcription factors, which further augments neurogenesis genes/proteins and enhances differentiation.

## MATERIALS & METHODS

### Cells

L-AN-5 cells were obtained from Children’s Oncology Group - COG Cell Line and Xenograft Repository (www.cogcell.org) and maintained in RPMI-1640 cell culture media (Sigma-Aldrich, St. Louis, Missouri, United States) supplemented with 10% fetal bovine serum (Gibco, 10082139) and 1% penicillin/streptomycin (Gibco, 15140122) at 37C and 5% CO_2_ for a maximum of 10 passages. To induce differentiation of L-AN-5 to a neuron-like phenotype, cells were incubated with RPMI/FBS supplemented with 10 μM retinoic acid (RA) (Sigma-Aldrich, St. Louis, Missouri, United States) for 4 days (media replaced every 2 days).

### Lentivirus vectors generation and transduction

For circZNF827 and hnRNPL knockdown we used dicer-independent small hairpin RNAs reported earlier to mediate efficient knockdown of circZNF827 in both the cytoplasm and the nucleus [1,11]. Third-generation lentiviral vectors carrying dishRNAs were constructed and produced in HEK293T cells as previously described [1,12]. All lentiviral preparations were made in at least triplicates and pooled before determination of viral titers. To determine viral titers of lentiviral preparations, flow cytometric measurements of eGFP expression were used as previously described [12]. eGFP expression levels were analyzed on a CytoFLEX flow cytometer (Beckman Coulter, Brea, California, United States). Lentiviral titers were calculated based on samples with between 1% and 20% eGFP-positive cells using the formula: Titer (*TU/ml)=F·Cn·DF/V*, where *F* represents the frequency of eGFP-positive cells, *Cn* the total number of target cells counted the day the transductions were carried out, and *DF* the dilution factor of the virus and V the volume of transducing inoculum.

One day prior to transduction with lentiviral vectors encoding dishRNAs, L-AN-5 cells were seeded at a density of 6.6 × 10^6^ cells per 10 cm dish. To achieve knockdown, cells were incubated with appropriate lentivirus dilution in RPMI media (3 MOI for each lentivirus, 6 MOI total) supplemented with 4 μg/ml polybrene (Sigma-Aldrich, St. Louis, Missouri, United States) for 24 hours, then media was replaced. Two days after transduction, L-AN-5 were subjected to differentiation by 10 μM RA (Sigma-Aldrich, St. Louis, Missouri, United States) for 4 days.

Following transduction and differentiation cells were harvested, counted, aliquots were taken to confirm knockdown, while the rest of the cells were fixed in 4% paraformaldehyde and immediately frozen. Cell aliquots were stored at −80C and used within one month of harvesting.

### PCR and RT-PCR

For RNA quantification and analysis, first-strand cDNA synthesis was carried out using Maxima H Minus cDNA Synthesis Master Mix, with dsDNase (Thermo Fisher Scientific, M1682) according to the manufacturer’s protocol. qPCR reactions were prepared using gene-specific primers (Supplementary table 3) and Platinum SYBR Green qPCR Supermix-UDG (Thermo Scientific) according to the manufacturer’s protocol. AriaMx Real-time PCR System (Agilent Technologies) was used for quantification. Relative RNA levels were normalized to GAPDH.

### CUT&RUN

CUT&RUN was performed using the CUT&RUN Assay Kit (Cell Signaling Technologies, #86652) according to the manufacturer’s protocol. Prior to footprinting, cell aliquots were thawed, counted and diluted to ensure 150,000 cells per reaction. The antibodies and dilutions used are listed in Supplementary table 4.

DNA fragments were purified from enriched chromatin samples using the DNA Purification Buffers and Spin Columns (ChIP, CUT&RUN, CUT&Tag) (Cell Signaling Technologies, #14209). Resulting ssDNA samples were stored at −80C and shipped on dry ice. Enrichment was confirmed using qPCR for RPL30 (CUT&RUN Assay Kit) and NGFR promoter (Supplementary figure 5).

DNA libraries were prepared, quality tested and sequenced at BGI on the DNBSEQ platform generating paired-end 100 bp reads. Low-quality and adapter sequences were removed from raw data using SOAPnuke [13]. Reads were aligned to the reference genome (GRCh38/hg38) using Bowtie2 v2.5.1 (*--local --very-sensitive --no-mixed --no-discordant --phred33 -I 10 -X 700*) (alignment statistics can be found in Supplementary figure 2 and table 5). Samples were normalized using a greenlist of genomic regions [14] and peaks were called against IgG enrichments using SEACR v1.3 [15] (*-non -relaxed*). Consensus peak sets were established, reads in peaks were counted from raw data with Diffbind [16] and used for differential enrichment analysis with DESEQ2 [17].

### RNA extraction and RNA-seq

To obtain RNA a portion of cells was taken when preparing CUT&RUN aliquots, dissolved in TRI Reagent (Sigma-Aldrich, St. Louis, Missouri, United States, 93289) and stored at −20C until extraction. Extraction was performed according to manufacturer’s instructions. RNA was dissolved in 20 uL nuclease-free water and split into aliquots stored at −80C.

The concentration and quality of RNA samples was assessed using Qubit and Bioanalyzer. Efficiency of circZNF827 knockdown was confirmed using RT-qPCR.

Samples were reverse transcribed and DNA libraries were prepared, tested and sequenced at BGI on the DNBSEQ platform generating paired-end 100 bp reads. Low-quality and adapter sequences were removed from raw data using SOAPnuke [13]. Reads were aligned to the reference genome (GRCh38/hg38) using STAR v2.7.11a [18]. Gene-level transcript abundance was quantified using featureCounts (Subread v2.0.6) [19] and used for differential expression analysis with DESEQ2 [17].

## Data handling and statistical analyses

Unless specified otherwise, data analysis and visualization were performed using R (version 4.5.1). Statistical tests were considered significant at p<0.05.

For transcription factor binding analysis, NR2F1 and POU4F1 binding motifs were obtained from JASPAR database [20] and FIMO from MEME suite [21] was used to calculate average scores for known human genes. GSEA was then used to test for enrichment of select gene sets among targets of either TF [22].

## Data and code availability

The data that support the findings of this study have been deposited in Gene Expression Omnibus *as GSE322550 (RNA) and GSE322552 (CUT&RUN*).

## Acknowledgements

This study was supported by the Independent Research Fund Denmark – Natural Sciences, case 026-00134B.

Data storage and high performance computing for this project were performed on the GenomeDK cluster. We would like to thank GenomeDK and Aarhus University for providing computational resources and support that contributed to these research results.

## Notes

https://www.ncbi.nlm.nih.gov/geo/query/acc.cgi?acc=GSE322552

https://www.ncbi.nlm.nih.gov/geo/query/acc.cgi?acc=GSE322550

### Competing Interest Statement

The authors have declared no competing interest.

### Summary of Updates

Supplementary figures updated (Figures S1-S7 were missing)

