## Supplementary figure for "Mechanism of circZNF827-mediated transcriptional repression during neuronal differentiation"

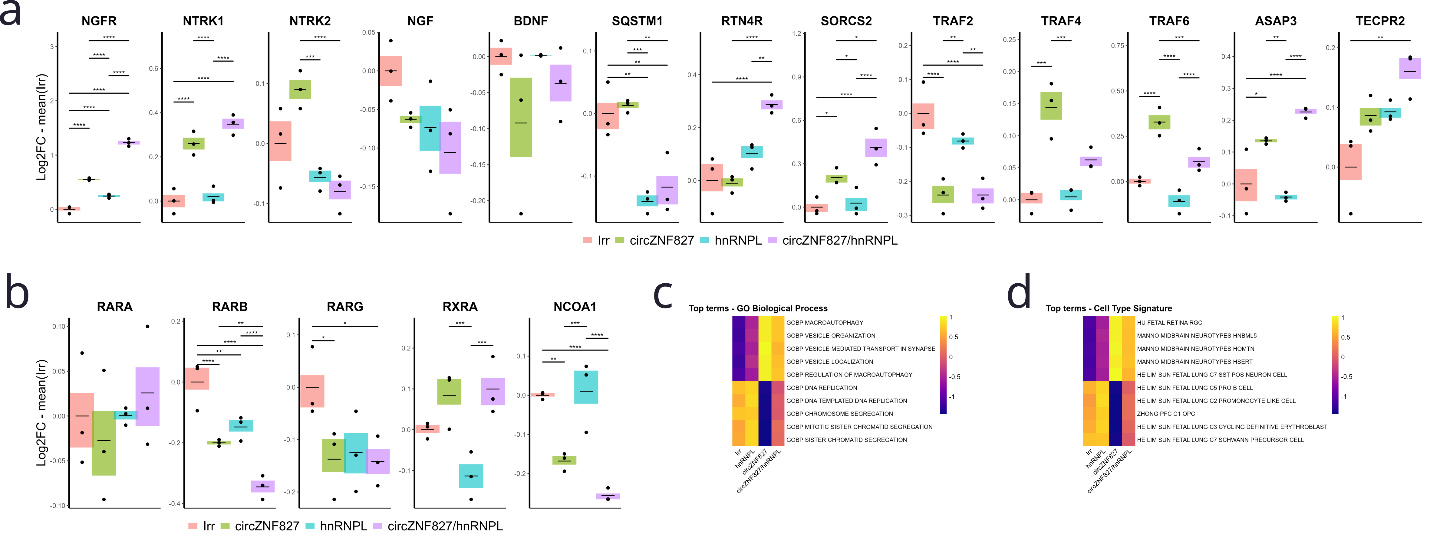

**Supplementary figure 1. RNA-Seq - additional plots.** (**a**) Boxplots of gene expression for genes in the NGFR pathway. Only genes with detectable peaks are shown. (**b**) Boxplots of gene expression for genes in the RAR pathway. (**c**) Heatmap of gene expression across the Gene Ontology Biological Process terms enriched in up- and downregulated differentially expressed genes. (**d**) Heatmap of gene expression across the Cell Type Signature (MSigDB) terms enriched in up- and downregulated differentially expressed genes.

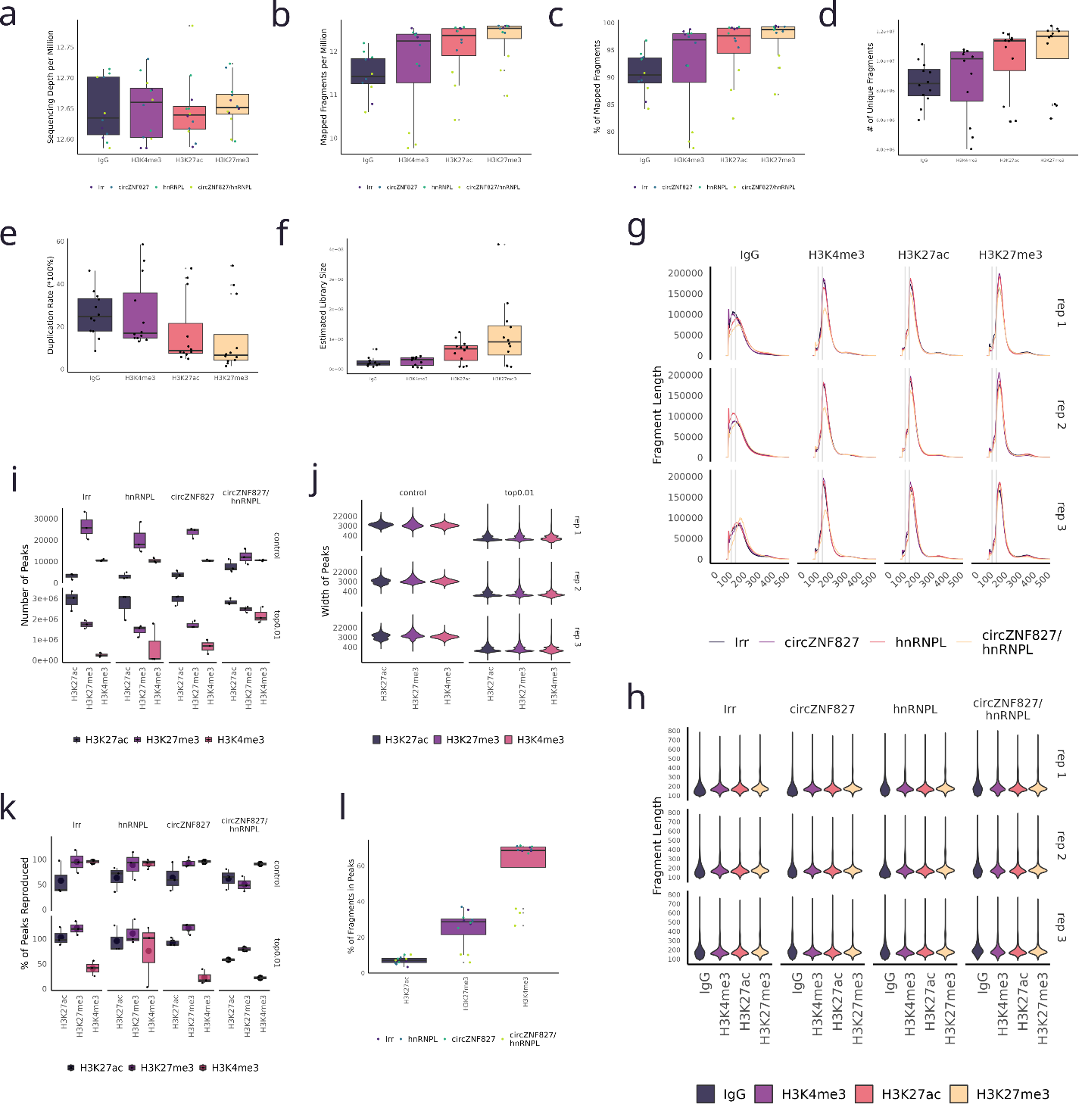

**Supplementary figure 2. CUT&RUN - read alignment and peak calling quality control plots.** (**a**) Sequencing depth per million bases. (**b**) Number of mapped fragments per million. (**c**) Percentage of mapped fragments. (**d**) Number of uniquely mapped fragments. (**e**) Fragment duplication rate. (**f**) Estimated library size. (**g,h**) Fragment length distribution by read length and by sample. (**I,j**) Number and size of peaks called by SEACR. (**k**) Reproduction rate of peaks. (**l**) Percentage of fragments in peaks (FriP).

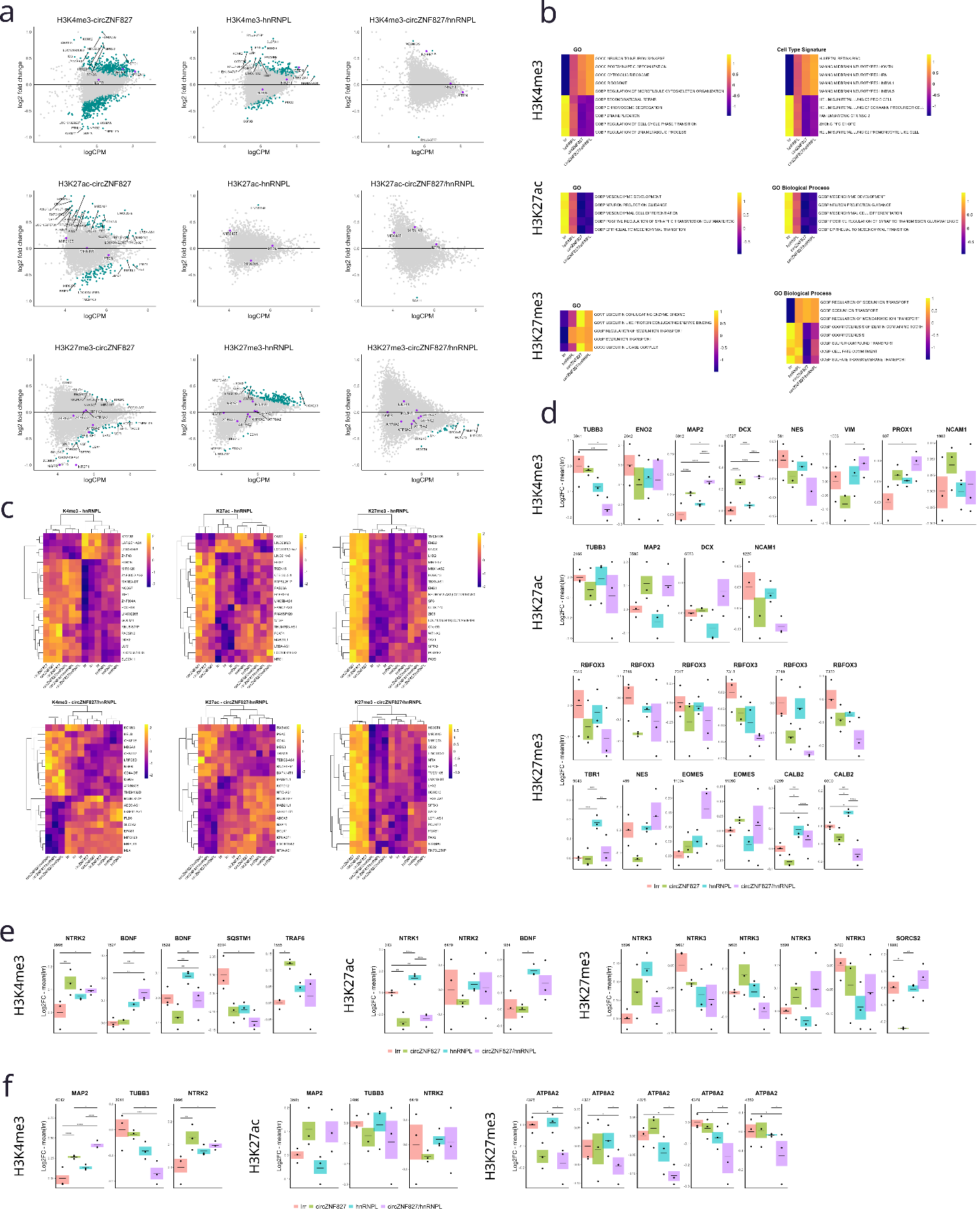

**Supplementary figure 3. CUT&RUN – additional analysis plots.** (**a**) MA plots of differentially enriched histone modification peaks for the three knockdown conditions. (**b**) Heatmaps of histone modification signatures across the Gene Ontology, Gene Ontology Biological Process and Cell Type Signature enriched in up- and downregulated differentially enriched histone modification peaks. (**c**) Heatmaps showing 20 top genes with differentially enriched peaks for the three histone modifications after circZNF827 knockdown. Only peaks in proximity to known genes are shown. (**d**) Boxplots showing histone modification levels for peaks associated with select neuronal markers. Only genes with detectable peaks are shown. (**e**) Boxplots showing histone modification levels for peaks associated with members of the NGFR pathway. Only genes with detectable peaks are shown. (**f**) Boxplots showing histone modification levels for peaks associated with select genes of interest based on previous data. Only genes with detectable peaks are shown.

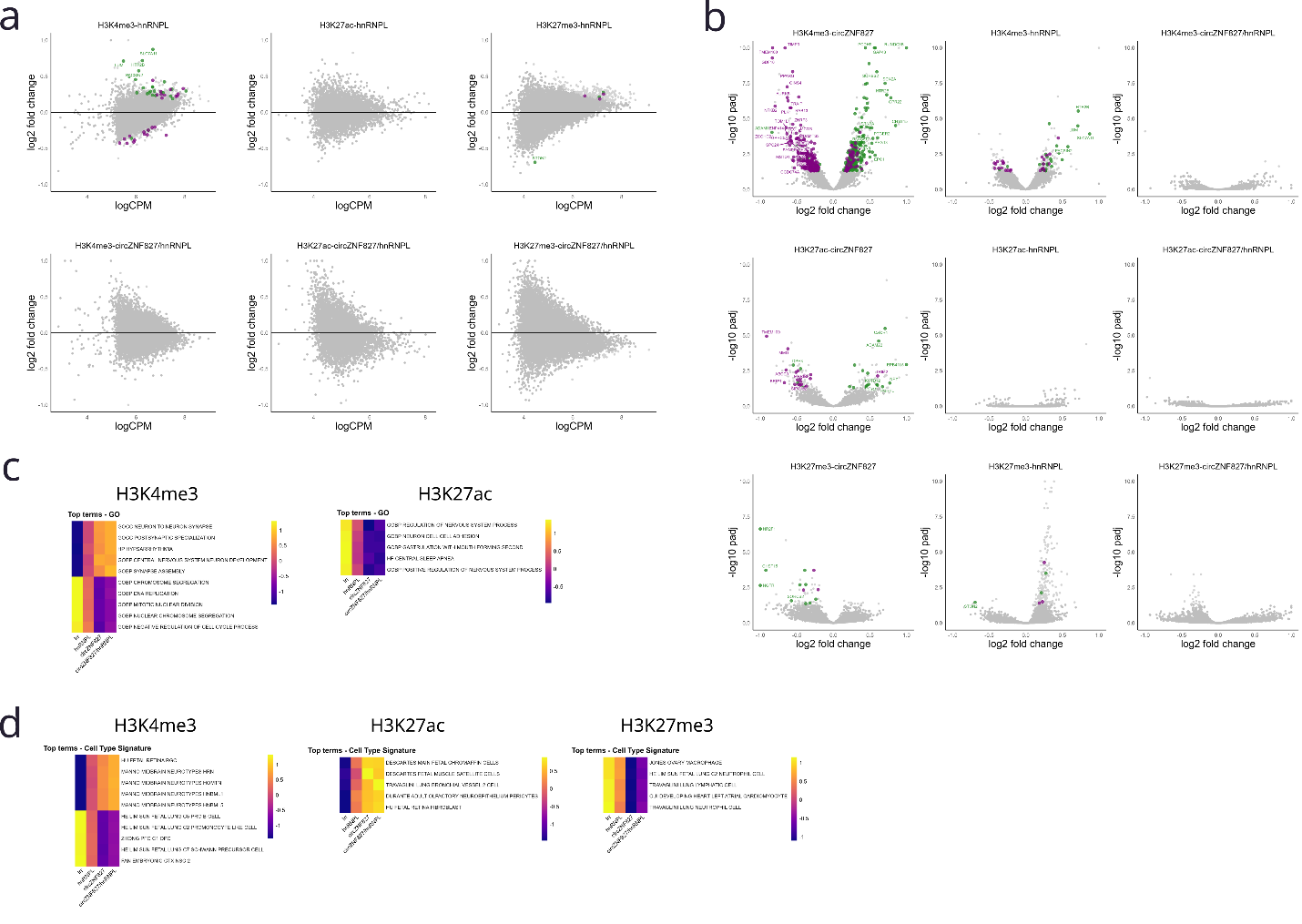

**Supplementary figure 4. Integrated omics analysis - additional plots.** (**a**) MA plots of differentially enriched histone modification peaks showing genes differentially expressed at the transcript level in green (up-regulated) or purple (down-regulated). (**b**) Volcano plots of differentially enriched peaks for all histone modifications showing genes differentially expressed at the transcript level in green (up-regulated) or purple (down-regulated). (**c**) Heatmaps of histone modification signatures across the Gene Ontology terms enriched in gene sets with concordant changes in histone modification and transcript abundances. (**d**) Heatmaps of histone modification signatures across the Cell Type Signature (MSigDB) terms enriched in gene sets with concordant changes in histone modification and transcript abundances.

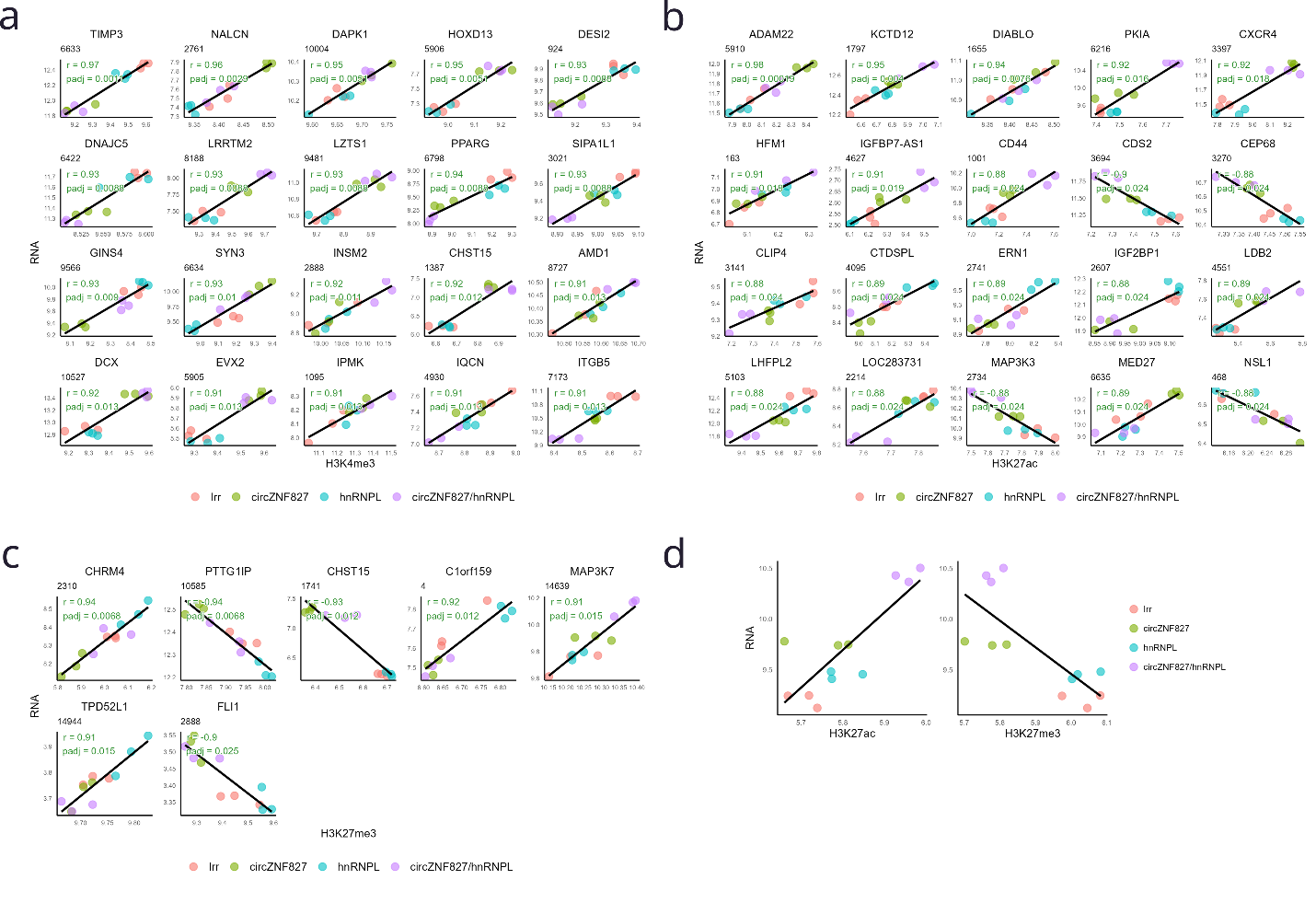

**Supplementary figure 5. Correlation of histone modification signatures and gene expression- additional plots.** (**a**) H3K4me3 levels and RNA expression for 20 genes with the highest correlation by adjusted p-value Benjamini-Hochberg). (**b** H3K27ac levels and RNA expression for 20 genes with the highest correlation by adjusted p-value Benjamini-Hochberg). (**c**) H3K27me3 levels and RNA expression for 7 genes with the highest correlation by adjusted p-value Benjamini-Hochberg). (**d**). Correlation between H3K27ac and H3K27me3 signal at the NGFR locus and its RNA abundance across all samples. Black solid lines indicate the general trend across all samples.

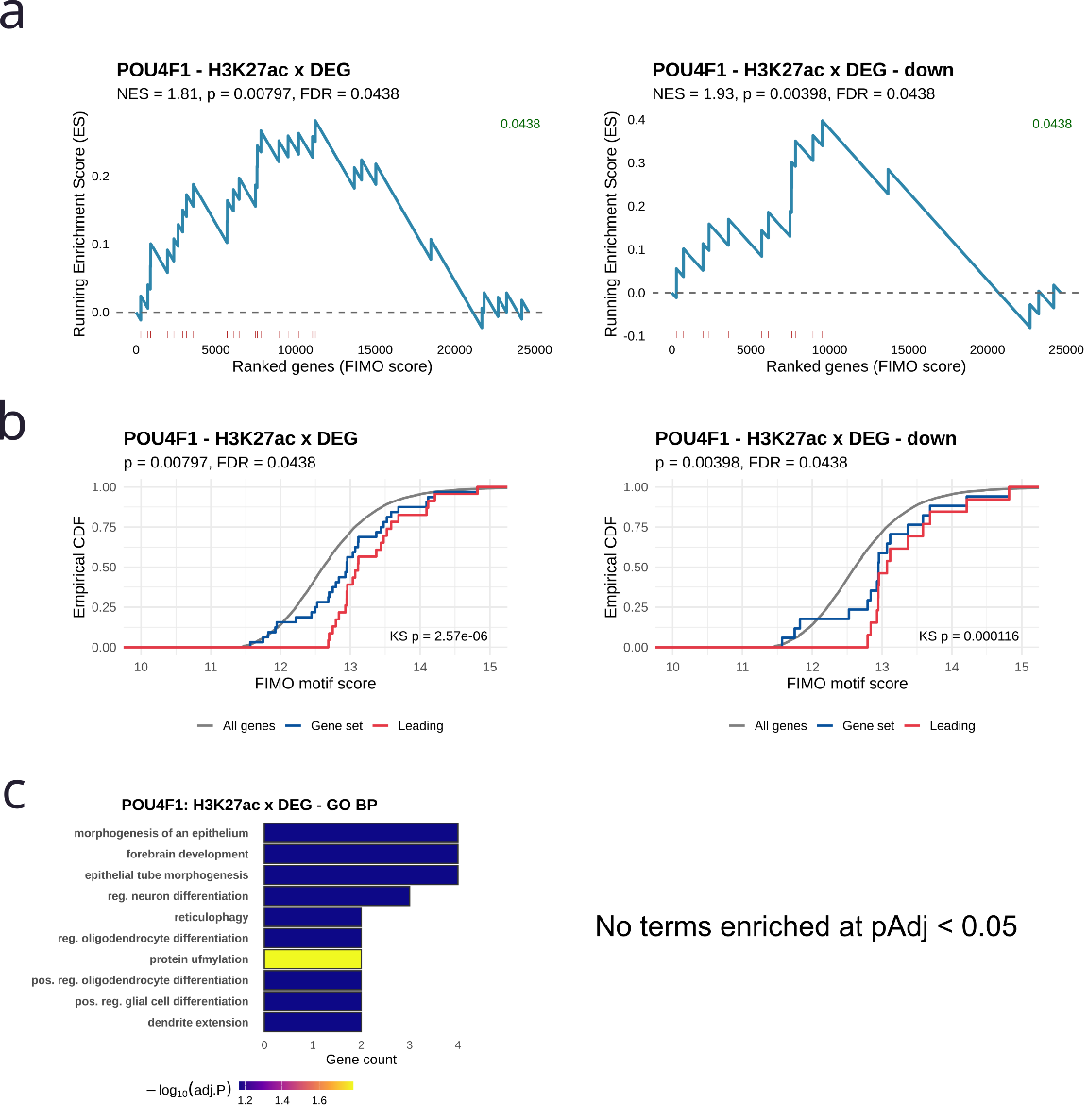

**Supplementary figure 6. Transcription factor enrichment plots - POU4F1.** (**a**) Gene Set Enrichment Analysis (GSEA) of gene sets based on CUT&RUN and/or RNA-Seq data against all predicted NR2F1 targets. (**b**) Empirical cumulative density function (CDF) plots of gene sets based on CUT&RUN and/or RNA-Seq data against all predicted NR2F1 targets. Kolmogorov-Smirnov test p-value (KS p) is given for comparison of Leading edge genes versus all ranked targets. (**c**) Gene Ontology Biological Process enrichments of Leading edge gene sets from the respective GSEA.

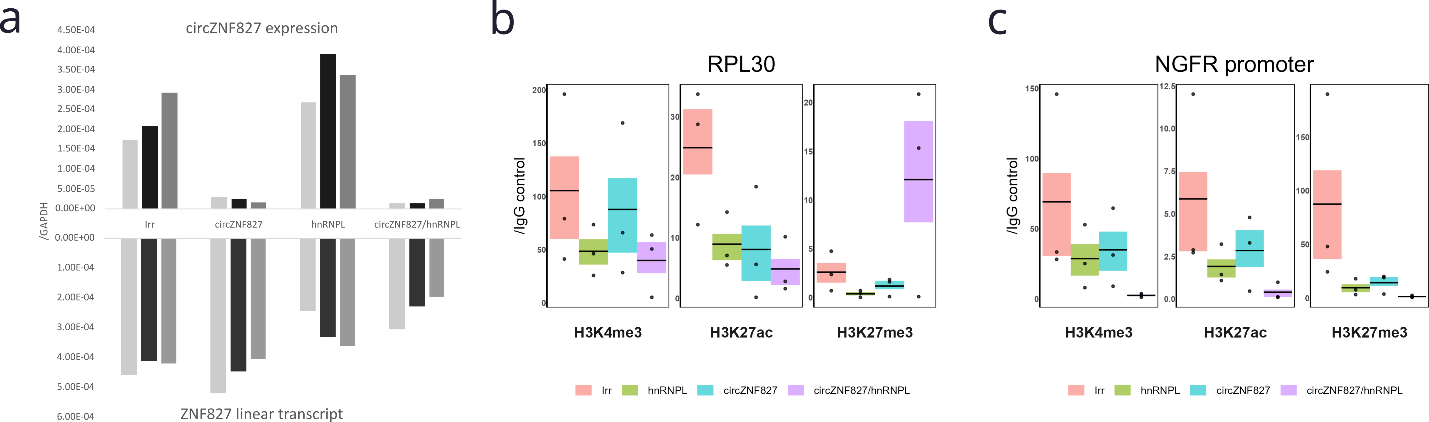

**Supplementary figure 7. CUT&RUN – experimental quality controls.** (**a**) Knockdown efficiencies for the circZNF827 and hnRNPL knockdowns. Expression levels were measured by qRT-PCR relative to GAPDH. (**b**) Chromatin enrichment efficiency for histone marks at the RPL30 locus. Enrichment levels were measured by qPCR relative to IgG control. (**c**) Chromatin enrichment efficiency for histone marks at NGFR promoter. Enrichment levels were measured by qPCR relative to IgG control.
